# JASS: Command Line and Web interface for the joint analysis of GWAS results

**DOI:** 10.1101/714832

**Authors:** Hanna Julienne, Pierre Lechat, Vincent Guillemot, Carla Lasry, Chunzi Yao, Vincent Laville, Bjarni Vilhjalmsson, Hervé Ménager, Hugues Aschard

## Abstract

Genome Wide Association Study (GWAS) has been the driving force for identifying association between genetic variants and human phenotypes. Thousands of GWAS summary statistics covering a broad range of human traits and diseases are now publicly available, and studies have demonstrated their utility for a range of secondary analyses. This includes in particular the joint analysis of multiple GWAS to identify new genetic variants missed by univariate screenings. However, although several methods have been proposed, there are very few large scale applications published so far because of challenges in implementing these methods on real data. Here, we present JASS (Joint Analysis of Summary Statistics), a polyvalent *Python* package that addresses this need. Our package solves all practical and computational barriers for large-scale multivariate analysis of GWAS summary statistics. This includes data cleaning and harmonization tools, an efficient algorithm for fast derivation of various joint statistics, an optimized data management process, and a web interface for exploration purposes. Benchmark analyses confirmed the strong performances of JASS. We also performed multiple real data analyses demonstrating the strong potential of JASS for the detection of new associated genetic variants across various scenarios. Our package is freely available at https://gitlab.pasteur.fr/statistical-genetics/jass.

## Introduction

The human genetics community has now access to a wealth of genome-wide association studies (GWAS) summary statistics for a wide spectrum of phenotypes, ranging from biometric measurements to molecular phenotypes and most common diseases. For example, as of May 2019, the NHGRI-EBI GWAS Catalog contains the results from 3989 GWAS^1^. The tremendous value and practical utility of summary data has been demonstrated for a range of secondary analyses^2–7^. Indeed, working with GWAS summary data solves both practical and ethical concerns, such as the protection of the anonymity of the participants, the secured storage of millions of variants across hundreds of thousands of individuals, as well as the harmonization of consortium data.

The joint analysis of multiple phenotypes based on GWAS summary statistics is currently a very active area of research with dozens of approaches published the past few years^8–16^. The primary goal of analyzing multiple phenotypes jointly is to increase statistical power to detect associated variants missed by univariate analysis^17–19^. However, large-scale real data multivariate GWAS analyses are still rare despite recent extensive effort from the community. While published methods have shown efficiency in simulated data and real data examples composed of few GWAS, we found more demanding applications, including dozens of GWAS performed from various platforms with partial sample overlap, to be very challenging. Among most prominent issues were: harmonizing heterogeneous summary statistics data format across studies, optimizing computation time to deliver joint analysis result in few minutes for several millions of SNPs, and summarizing and comparing joint analysis results versus univariate results.

Moreover, an important aspect of exploring complex multivariate data –here GWAS summary statistics of multiple phenotypes–is the visualization and management of results and data. This is a common issue in human genetics, and a number of software have been published to help investigators coping with this issue. There are now many user-friendly tools allowing for the annotation of results based on existing functional database (e.g. Dalliance^20^, toppar^21^), the secure storage and extraction of results from database (e.g. gwas ATLAS^7^), or the plotting of results integrating specific features (e.g. LocusZoom^22^, Assocplot^23^). Similarly, the dissemination of the methodology for the joint analysis of multiple GWAS summary statistics, requires the development of robust and computationally efficient tools. With thousands of GWAS studies now publicly available, this requirement has become even more crucial.

Here, we present JASS *(Joint **A**nalysis of **S**ummary **S**tatistics),* an integrated package for the joint analysis of multiple GWAS summary statistics, including at the same time analytical tools, visualization functions, and an embedded web-interface. More precisely, JASS provides the following features: 1) functions for the fast and efficient computation of joint statistics from dozens to hundreds of GWAS results, 2) an interactive web server for the visualization of a selection of GWAS, along with the result of their joint analysis, 3) the possibility to install the software locally, so that interested users might apply the analysis to their own data in the safety of their own computation facility, and 4) a command line interface for advanced users. The paper is structured as follows: first we detail the main functionalities and the visualization tools of JASS; second, we discuss the performance optimization strategies we adopted; third, we provide some technical details of the package; and last, we present applications using real GWAS summary data performed using the public JASS server.

## Material and Methods

### Overview of JASS

JASS is a *Python* package that handles the computation of joint statistics over sets of selected GWAS results through either a command line interface and or a web interface. The derivation of joint statistics, and the generation of static plots to display the results, as well as more advanced features such as the implementation of user-defined statistics, can be easily performed using the command line interface. Many of these functionalities can also be accessed through a web application embedded in the *Python* package, which also enables the exploration of the results through a dynamic *JavaScript* interface. The list of available functions and features are provided in **Table S1** and **S2. Figure 1** shows a standard command line analysis workflow including the steps performed through companion packages.

### Statistical tests implemented in JASS

A number of methods have been published for the joint analysis of multiple GWAS summary statistics. We implemented two approaches in the current version that cover most of the existing joint tests, although users also have the possibility to implement their own joint statistic and plug it in the analysis. The first test is a standard omnibus approach which combines *k* single GWAS statistics to form *k* degrees of freedom (df) statistics. For a given SNP, the omnibus statistic can be expressed as:

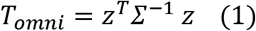

where *z* = (*z*_1_,*z*_2_, …, *z_k_*) is a vector of *k Z*-scores, derived from the available GWAS summary statistics as 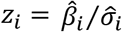 where 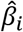 and 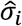 are the estimated regression coefficient and its standard error for study *i*; *Σ* is the covariance matrix between the *Z*-scores and is assumed unique for all SNPs analyzed. Under the null hypothesis of no association between the SNP tested and any of the *k* phenotypes tested jointly, *T_omni_* follows a *χ*^2^ distribution with *k* degrees of freedom.

The second statistic is a weighted sum of Z-scores and has the following form:

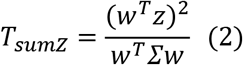

where *w* is a vector of weights assigned to each GWAS, and applied uniformly to all SNPs analyzed. Under the null hypothesis of no association between the SNP tested and any of the *k* phenotypes tested, *T_sumZ_* follows a *χ*^2^ distribution with 1 degree of freedom. This joint test is the most discussed one in the literature, and variations of this approach mostly consist in optimizing the weighting scheme^10, 11^.

Note that whichever multivariate statistic is used, the estimation of *Σ* is critical to ensure a correct type I error rate. As done in our real data application, we strongly recommand building that matrix using the intercept derived from univariate and bivariate *LDscore* regression described by Bulik-Sullivan et al^6^. However, users can provide their own estimation of the covariance matrix or use a default estimator provided by JASS.

### JASS input and output

The JASS package requires for input a set of GWAS summary statistics, and some metadata about these GWAS (e.g. coded allele, sample size per SNP...). Two additional key arguments can be provided: i) a matrix of the covariance between statistics under the null hypothesis (i.e. *Σ,* see equation (1) and (2)), and ii) a map of regions that is used to create summary results. If the covariance matrix is not provided by the user, it will be derived based on the observed pairwise correlation between GWAS after filtering out SNPs with *p*-values below a significance threshold (*P* < 5×10^−5^), in order to remove likely associated variants^24^. The primary purpose of the region map file is to improve exploration and visualization of the results. For meaningful interpretation of the region-based results, we strongly suggest to define the regions based on the SNPs Linkage Disequilibrium (LD) between variants. If not provided by the user, JASS will use regions based on LD recombination hotspot computed using the approach proposed by Berisa and Pickrell^25^ for European populations. From this input data, the command line tool allows for multiple joint tests to be performed: i) the Omnibus approach (equation (1)), ii) the sumZ approach (equation (2)), where the user has to specify a vector of weights, and iii) any other alternative statistics applicable to a vector of *Z*-scores.

Two main static plots can then be generated from the joint analysis: 1) a so-called *Manhattan plot,* i.e. a scatter plot of genome-wide association signal showing the −log_10_ of the *p*-value of each region, according to their position on the genome, and 2) a *Quadrant plot,* which allows for a fast comparison of *p*-values between the joint test and single GWAS results. In brief, the −log10 of the minimum *p*-value of each region from the joint analysis is plotted as a function of the −log10 of the minimum *p*-value from univariate GWAS analysis. The results is a 4 quadrants scatterplot where: (i) the upper left quadrant contains all the regions where the joint analysis is significant when all the original GWAS are not, hence newly detected regions; (ii) the upper right quadrant contains all the significant regions identified with both strategies; (iii) the lower right quadrant contains all the regions that are significant for any one of the selected GWAS but not for the joint analysis; (iv) and finally, the lower left quadrant displays the regions that contain no signals.

The web interface provides a complementary set of tools for both analysis and visualization. Note that it is applicable after all input data have been harmonized and merged along all required information (**Figure 1**), and it currently allows only for the omnibus test to be performed. Once a set of GWAS has been analyzed, the user can proceed to the exploration at the SNP level by clicking on a region. This action will trigger the representation of a SNP level heatmap of the Z-score across all GWAS analyzed jointly and a zoomed Manhattan plot of the joint test association results. When implemented on a public server, the web interface also offers the possibility of sharing the results from an analysis through a “Share direct link” button. It allows the user to generate a unique link for the joint analysis they performed, avoiding other investigators to replay the same analysis multiple times and an easy way to access additional details from a published study.

**Figure 1.**
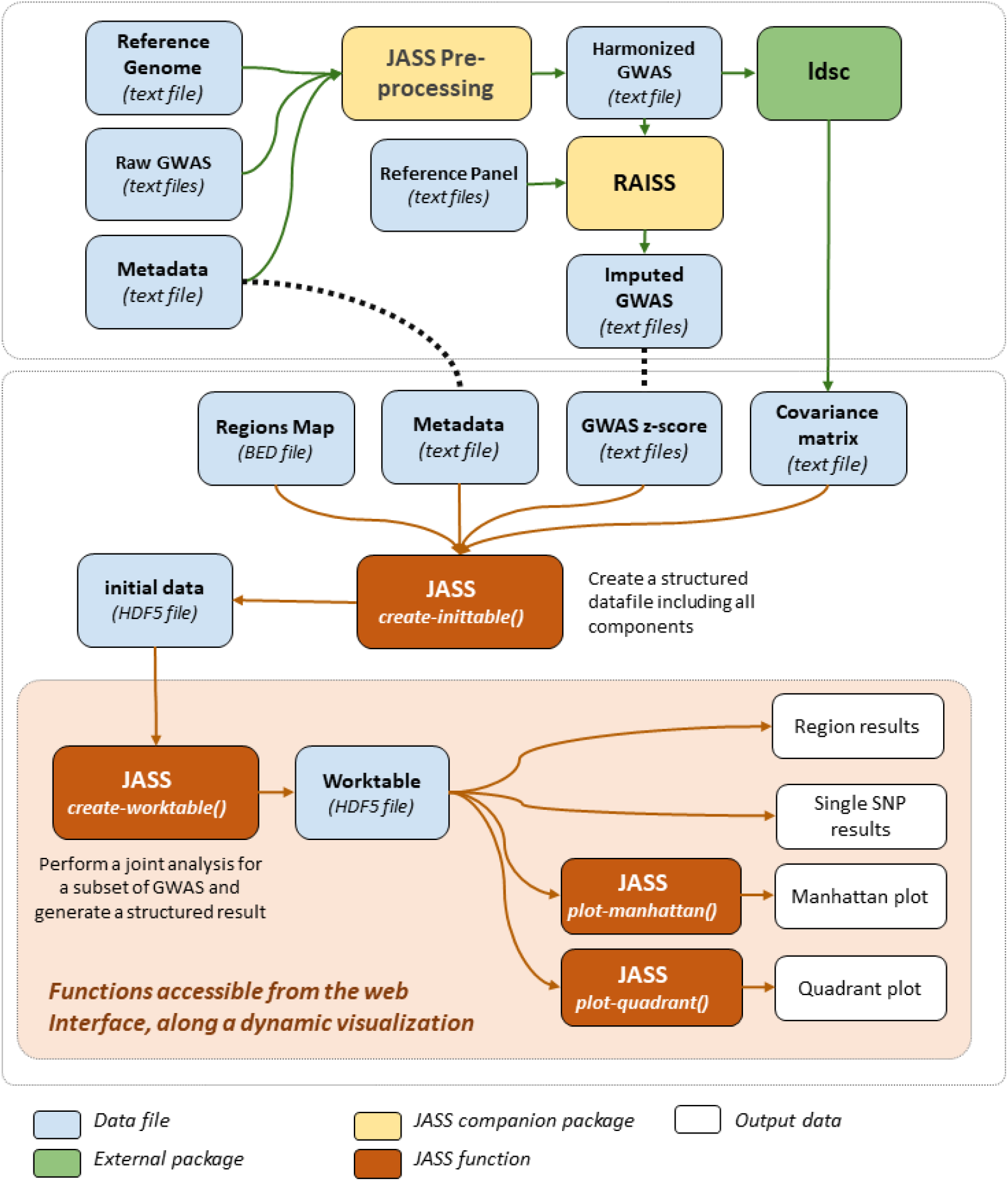
JASS workflow. The figure below shows the main steps of an analysis with JASS, including the pre-processing steps performed using either companion packages (in yellow) of external software (in green).

### Data pre- and post-processing

A critical issue to ensure a valid multi-GWAS analysis is the harmonization and cleaning of the input data. Raw GWAS data are usually heterogeneous in their content and format. To avoid the user series of time-consuming operations, we automatized all of these steps into a pre-processing *Python* package tool (jass_preprocessing) provided on behalf of JASS. This companion package addresses the following: i) it maps heterogeneous column names to standard names (rsID, pos, A0, A1, Z); ii) it derives Z-scores from *p*-values and signed statistics (log odds ratio or linear regression coefficients); iii) when missing, it infers per-SNP sample size from the effect estimate standard deviation and the minor allele frequency; iv) it filters SNPs with sample size lower than 75% of the maximum; and v) it aligns coded genetic variants with a reference panel. This reference panel should be filtered out for strand-ambiguous variants and variants with low frequency. In our example we used 6,978,319 SNPs selected using the European ancestry samples from the 1KG project^26^. This reference panel is available on the preprocessing package <u>gitlab</u>.

Importantly, joint statistics commonly require complete data to perform a statistical test. To increase the number of SNPs with complete data, we chose to perform the systematic imputation of GWAS included in our real data examples. Because of the complexity of the task, we developed a tool for the imputation of missing statistics in an independent study implemented in the <u>RAISS</u> package^27^. The input and output format of *RAISS* are the same as input format for the JASS package so the imputation step can easily be integrated in a JASS pipeline. Nevertheless, the imputation of missing data can only be partial, and therefore, JASS includes a computationally optimized procedure to derive joint statistics for each SNP only from the subset of GWAS with available data.

Finally, despite carefull pre-processing, it remains possible that some features susceptible to impact the validaty of multivariate tests are missing from the input data. For example, GWAS might include summary results for imputed SNP with relatively low imputation quality score (e.g. 0.3 ≤ *R*^2^ ≤ 0.7). Such variants do not impact the validity of the univariate screening, but can harbor differences in their covariance *Σ* as compared to the other SNPs. Such differences can induce false signals in a multivariate test. To handle such situations, we also propose a post-hoc filtering to remove likely outlier signal. It consists in removing SNPs with genome-wide significant multivariate signal (i.e. *p*-value belows 5×10^−8^) in the absence of other signal with a *p*-value belows a user specified threshold (10^−6^ by default) in the same region.

### Technical details

#### Architecture

JASS is a *Python* package that can be run both as a command line tool, and as an embedded web application, either on a local or a public web server. An overview of its architecture is illustrated in **Figure 2**, that describes its primary components.

**Figure 2.**
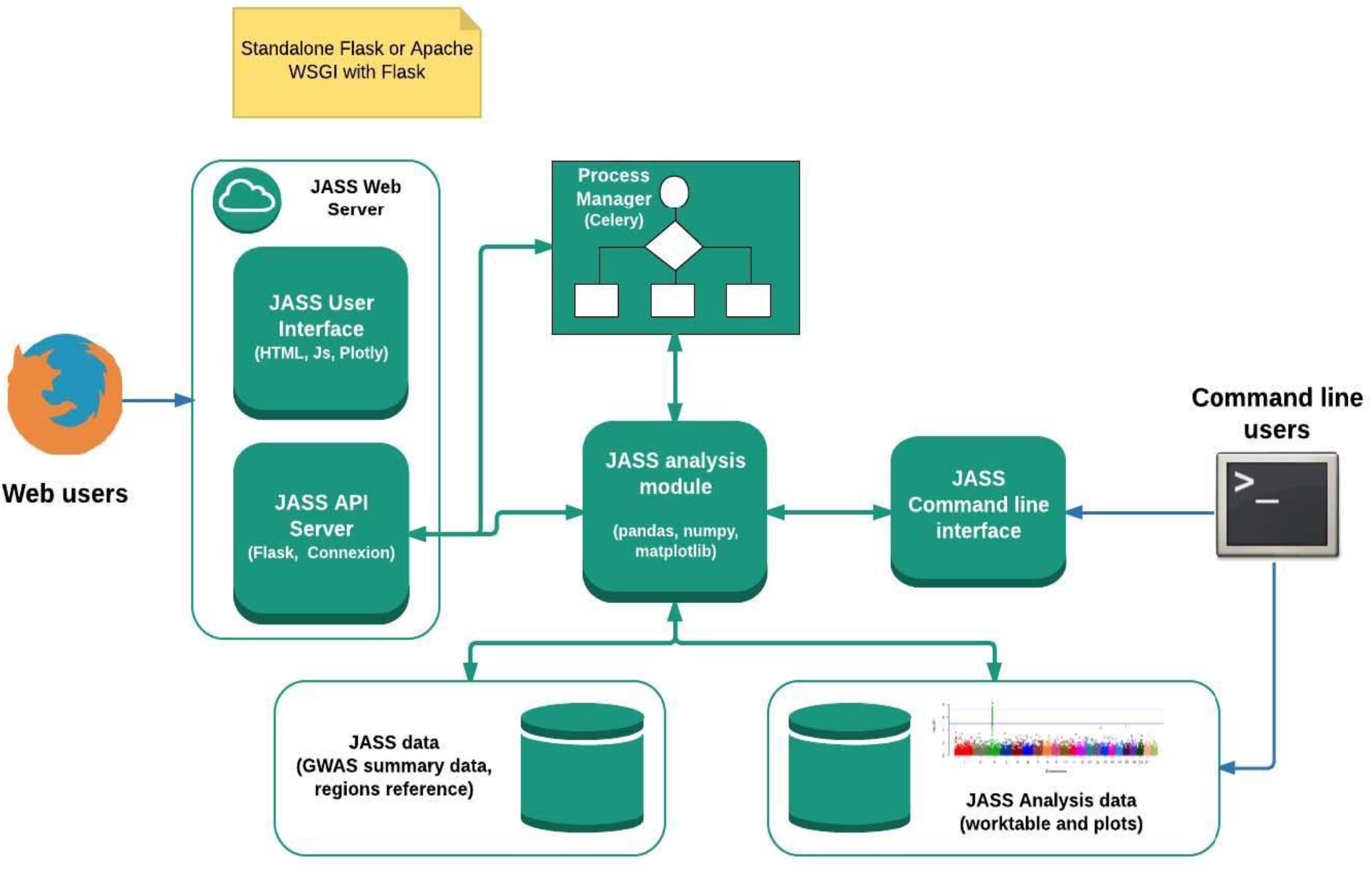
JASS architecture. The JASS *Python* package is composed of a central “analysis module” that defines computational features, enables reading input data and writing results to HDF files, and generating static plots as PNG files. These features can be accessed from a command line interface, or from an embedded web application powered by a Flask server that uses the connexion library and a *JavaScript* user interface using the plotly library that enables the execution of joint analysis and the exploration of the results. In the latter case, the management of the analysis jobs requires the setup of a Celery server.

#### Data storage and computations

The analysis module of JASS uses the *pandas* library for data processing, allowing for convenient and fast computations over large datasets. Furthermore, *pandas* includes a native support for *HDF5* files, which JASS uses to store both the initial data and the results of joint analyses. This format presents multiple advantages, including (1) generating cross-platform and compressed files, (2) enabling indexing (which later reduces the access time when accessing the results during the interactive exploration) and (3) storing multiple data frames in a single file.

#### Interactive web interface

The JASS web interface is implemented in *JavaScript, HTML* and *SVG*. To browse the results efficiently, we use a *plotly* library. This library enables the simultaneous display of very large numbers of points, as encountered in JASS, where the size of GWAS matrices to be explored can potentially reach hundred phenotypes per several million SNPs. It is already used in other GWAS analysis R packages, including for example <u>Manhattanly</u>. The tables we use for selecting phenotypes and exporting results use the jquery <u>datatable</u> plugin.

#### Web server

The web server provides all functionalities (except for the initial data import), through a *RESTful* API. This API is described with *OpenAPI,* and connected to the *Python* JASS analysis module through the *connexion* library. It is used primarily by the web user interface, but could also be used to run JASS analyses remotely, or with other client frameworks. Contrarily to the command line API, the statistics computation and plots generation operation is launched asynchronously, to avoid the potential timeouts that can occur during its execution. The server uses the *Celery task queue* to run these operations.

## Results

### Balancing resource usage

The amount of data that has to be managed and processed guided us to choose the architecture and the output format for JASS. As discussed in the previous sections, there are currently more than 3,000 GWAS summary statistics available. These GWAS currently contains approximately between 300K and 10 million genetic variants. However, future studies including either additional imputed SNPs, or sequence data, can potentially bring this total to up to 100 million variants or more. Given these numbers, the loading and processing of a multi-GWAS analysis would be intractable when using either a standard laptop or a standard web server (which includes often less than 10 Gigabytes of RAM). Therefore, emphasis has been put on optimizing the memory usage, while preserving reasonable execution times. This translated into two main components in the JASS implementation.

First, while the analysis is done on a single SNP basis, we constructed a region-based result on top of the single SNP results. In brief, we defined genomic regions based on the SNPs linkage disequilibrium (LD), and derived statistics and summary based only the most significant SNP from each region. This region-based result allows for both a very fast visualization at the genome level in the web interface and a preliminary count of independent signals, circumventing the loading of the whole genome results. Here we propose using regions based on LD recombination hotspots computed using the approach proposed by Berisa and Pickrell^25^. Note that JASS offers the possibility of using an alternative user-defined region map. The map we used in our examples includes 1,703 regions. We derived in **Figure 3A** the distribution of the number of SNPs per region in three arbitrary scenarios: 1M SNPs from the *Illumina Omni1-QuAd;* 9.9M SNPs after imputation as available in^28^; and 96M SNPs, as available in the latest UK Biobank genotype panel after imputation^29^. Overall, our approach shows a good scalability with an average of 590, 5812, and 54,664 SNPs per region, with maximum of 3,470, 19,217, and 145,987 SNPs, respectively.

**Figure 3:**
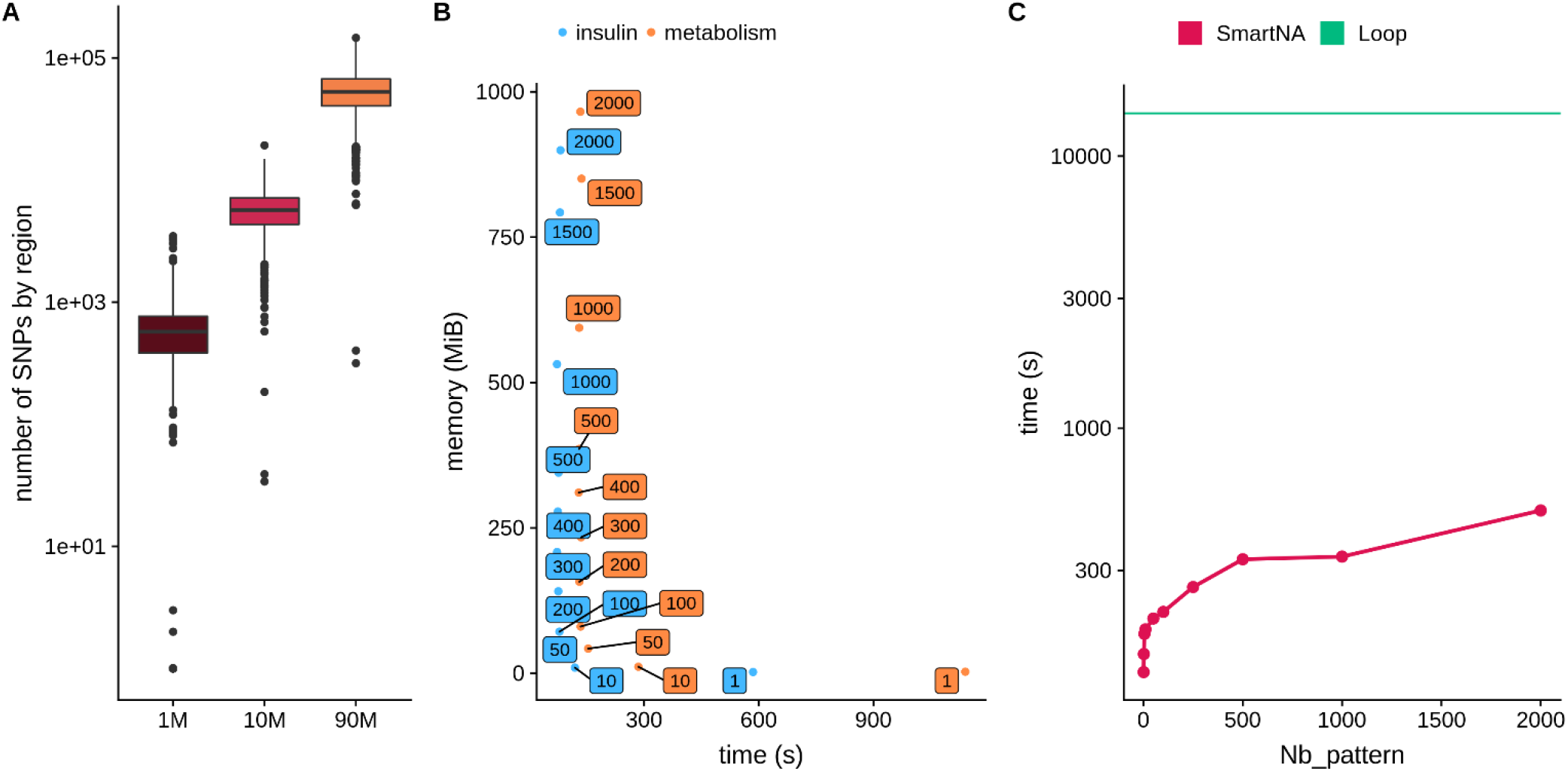
Resource usage and computation time. Panel **A)** shows the distribution of the number of SNPs per region when using the recombination hotspot map proposed by Berisa et al. 2016 while using 1M SNPs from the Illumina Omni1-QuAd; 9.9M SNPs from a recent GWAS; and 96M SNPs, as available in the latest UK Biobank genotype panel after imputation. Panel **B)** shows the trade-off between memory usage and computation time. JASS omnibus statistics was computed for 4 traits and 3,192,045 SNPs (insulin) and 11 traits and 3,673,285 SNPs (metabolism). The y-axis is the peak in memory usage in megabytes and x-axis the time (in seconds) to perform the task. Labels attached to data point give the number of genomic regions processed at once. Color represents the group of traits. Panel **C)** shows the computation time optimization. We computed the omnibus statistics for 3,673,285 SNPs for 11 traits, the y-axis is the time in seconds (logarithmic scale) and x-axis is the number of missing data in a pattern. The straight green line shows the result from a naïve computation loop approach over all SNPs, and the red line shows the results from our optimized algorithm.

Second, we implemented a strategy where the computation of the joint analysis is performed iteratively on chunks of 50 regions. The choice to process 50 regions at a time corresponds to an optimal trade-off between computation time and memory usage when analyzing multiple phenotypes (e.g. 20 or more, **Figure 3B**). Processing less regions would increase computation time and processing more regions would increase dramatically memory usage. The resulting performance makes it possible to compute a joint analysis in less than a minute, while preserving the stability of the server. This process is particularly critical when using the JASS webserver. Additionally, note that the *Celery* server is by default configured to queue and run sequentially the analyses launched potentially simultaneously on different datasets, to avoid excessive loads on the memory or CPU usage of the server. Finally, when analyzing jointly a large number of phenotypes (e.g. more than a hundred), command line on a local instance of JASS should be preferred.

### Optimized algorithm for fast computation

It is well established that computation time of genome-wide screening can be decreased dramatically by using matrix product rather than through an iterative loop (e.g.^30^). For illustration purposes, we considered an example of a real GWAS dataset including 11 traits and 3,673,285 SNPs after filtering out SNPs with missing data. The naïve loop-based implementation (computing the statistic for each SNP one-by-one and inverting the *Σ* matrix each time) took nearly 4 hours. On the other hand, processing the same data with a matrix product took 127.1 seconds (**Figure 3C**). However, matrix-based derivation cannot be readily implemented in the presence of missing data. Missing values imply that the statistics will differ with rows according to which traits are missing. This is a particularly acute problem for the omnibus test, which requires the inversion of the covariance matrix for each SNPs.

To solve this issue, we implemented an optimized algorithm that treats jointly genetic variants displaying the same missing value pattern. The computational cost of this operation can be factorized because values are generally not missing at random: missingness depends on the coverage of the genome by the technology used for the studies, the technology itself, and the genotype imputation approach that was used. The algorithm first identifies patterns of missing values in the Z-scores matrix, and computes *Σ^−1^* only once for each pattern. Each set of SNPs harboring the same pattern is then analyzed using a matrix-based derivation. **Figure 3C** reports the execution time as a function of the number of missing values patterns in the data using the aforementioned real data example, after randomly incorporating an increasing amount of missing data. As shown in this figure, even with the number of missing values patterns set at its theoretical maximum (for 11 traits, this would be 2^11^ = 2,048 patterns), the execution time of the matrix-based computation is dramatically lower than the execution time of the naïve loop implementation (t=499s *versus* t=14,365s, respectively).

### Comparison to the existing code to perform joint analysis

We compared the functionalities and computational performances of JASS against other approaches for the joint analysis of GWAS summary statistics. Out of the eight studies (**Table S3**), two did not provide code^14, 31^, two provided loose R functions^11, 12^, one provided *Python* 2.7 scripts^3^, one provided code in a non-open source software^9^, and two provided R packages^10, 13^. The quality of documentation varied greatly from one method to another going from its absence to a wiki describing in depth routine usage. Only one package offered a command line interface. None of the advanced features offered in JASS, such as the management of missing values, an accompanying *Python* 3 package to harmonize and clean heterogeneous GWAS summary statistics, and the possibility to deploy an interface server, were available in other approaches.

Finally, to compared the execution time of JASS with the HIPO^10^ and MTAG^3^ methods on a lipid example containing 4 traits and 1,818,293 SNPs. HIPO, MTAG and JASS ran respectively in 329, 212 and 33 seconds. To be fair with the HIPO methods the reported running time includes the estimation of the genetic correlation matrix by the LD score. Note that no option is provided in the function to avoid repeating the estimation for each analysis.

### Detection of new associations with JASS

We deployed an online implementation of JASS which currently includes 49 publicly available GWAS summary statistics (http://jass.pasteur.fr/index.html, **Table S4**). These GWAS cover several common diseases (e.g. asthma, type 2 diabetes, cardiovascular diseases) and quantitative traits (e.g. body mass index, total cholesterol). All GWAS were pre-processed using the JASS companion package and were aligned to a reference panel of European ancestry sample from the 1KG Project Phase 3 data^26^. Rare (MAF < 1%), non-biallelic and strand ambiguous SNPs were filtered out from the reference panel. These 49 GWAS represent only a fraction of all publicly GWAS available, however there are already 5.6×10^14^ possible combinations of phenotypes that can be analyzed jointly from this subset. As there is currently no established strategy to determine which specific set of phenotypes should be considered for joint test when addressing a specific biological question, it illustrates the strong need of disseminating to the community a fast and user-friendly tool for multi-GWAS analysis. Our package offers the possibility of performing multiple exploratory analyses extremely fast. To illustrate and demonstrate the potential of JASS, we performed three real data applications using these data.

#### Example 1

In this first example, we considered a simple scenario where an investigator is interested in complementing results from genome-wide genetic association studies of multiple insulin phenotypes. The goal is to perform a multivariate analysis to identify additional genetic variants associated with insulin phenotypes missed by the univariate analyses. Here, we performed the analysis using the online version of JASS, and as mentioned in the method section, we generated a unique and publicly available link for this specific analysis <u>here</u>. We ran the omnibus test for the 4 insulin phenotypes from the MAGIC consortium (insulin resistance, fasting insulin, Insulin secretion, and fasting proinsulin) available (**Table S4**). The main steps and corresponding screenshots of the analysis are presented in **Figure 4**. After imputation and filtering SNPs with data on at least two phenotypes, a total of 1,657,972 SNPs was available for the analysis.

**Figure 4.**
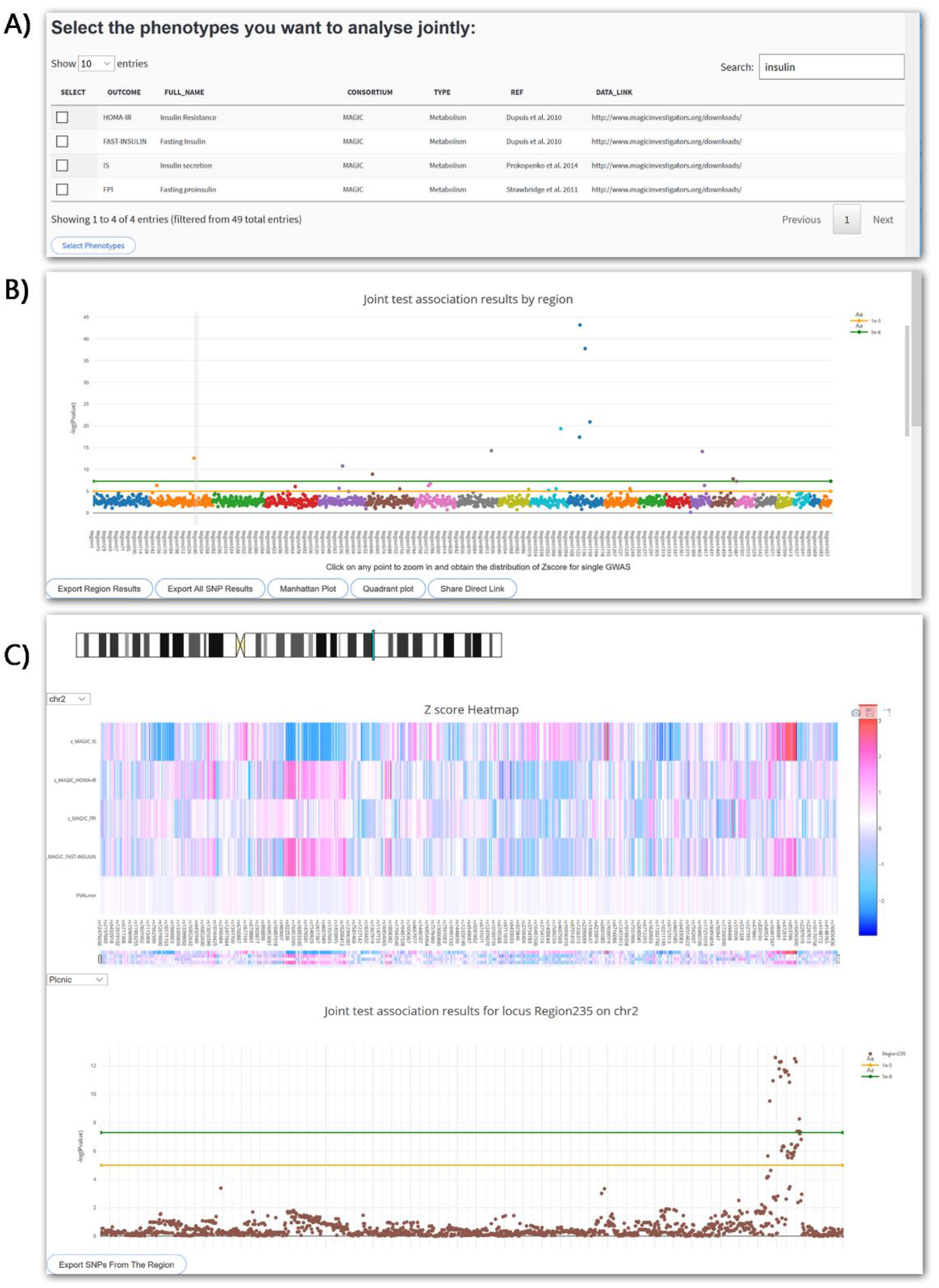
Screenshot JASS interface for example 1. We present below screenshots from the web interface of the 3 main steps for the analysis performed in application 1. A) Insulin-related GWAS available in the database are selected (insulin resistance, fasting insulin, Insulin secretion, and fasting proinsulin), B) the genome wide results from the joint test are presented in a Manhattan-like plot of the omnibus test, showing the top −log10(P) per genomic region, C) when a region of interest is selected, a dynamic heatmap of the Z-scores of the previously selected phenotypes is generated along a zoom on the Manhattan plot showing the −log10(P) single SNP signal for the joint analysis.

The univariate analyses identified 11 regions of interest (**Table 1**). All except two were also significant with the joint test. Conversely, the omnibus approach identified one new region in chromosome 2 within the G6PC2/ABCB11 locus. The top variant from the omnibus test for that region was rs560887 (P=2.6×10^−13^). None of the *p*-value for the univariate test was nominally significant for that variant (P equal 0.44, 0.10, 0.11, and 0.52 for insulin resistance, fasting insulin, Insulin secretion and fasting proinsulin, respectively). However, rs560887 has been reported to be associated with multiple phenotypes, including in particular elevated fasting plasma glucose and increased insulin release after oral and intravenous glucose loads^32^. The difference in significance between univariate and multivariate association suggests a genetic effect on a combination of the original phenotypes (e.g. difference or ratio between 2 insulin traits).

**Table 1:**
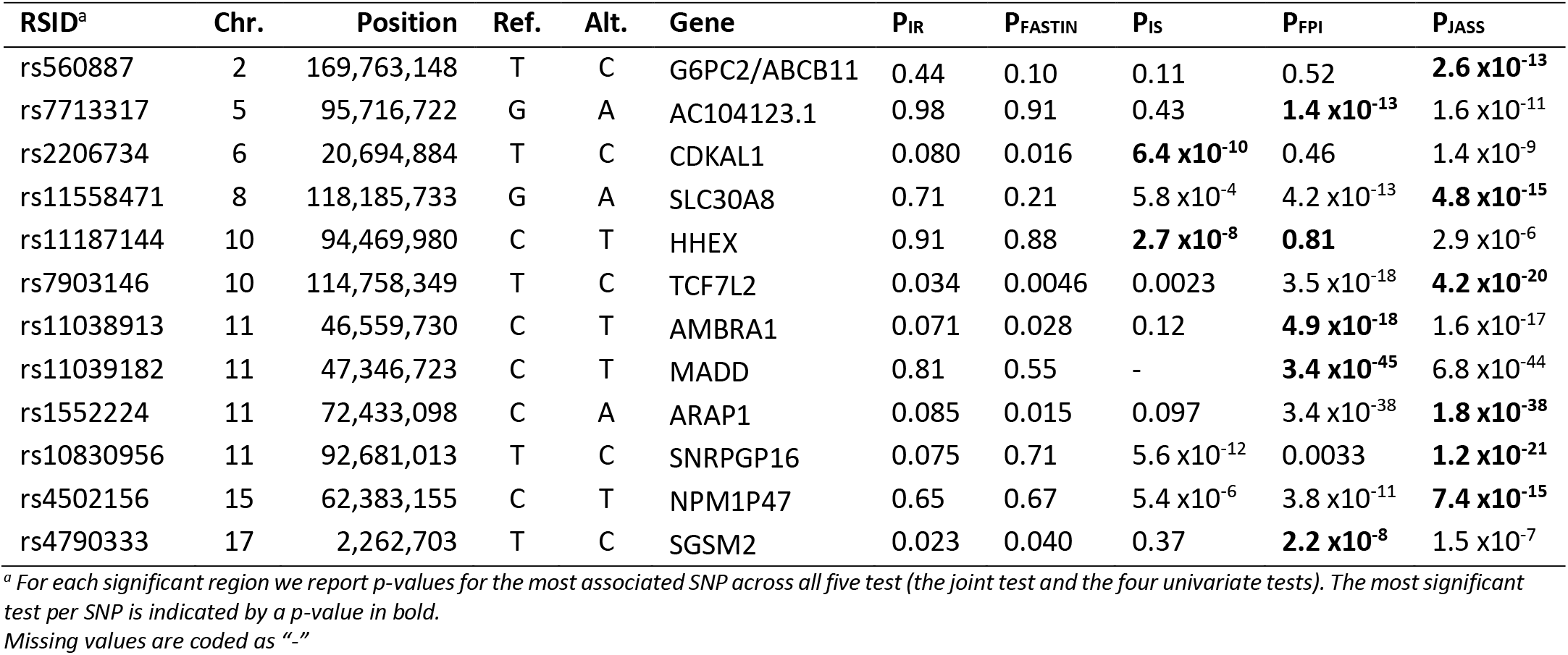
Top associated SNP from example 1

| RSID<sup>a</sup> | Chr. | Position | Ref. | Alt. | Gene | P<sub>IR</sub> | P<sub>FASTIN</sub> | P<sub>IS</sub> | P<sub>FPI</sub> | P<sub>JASS</sub> |
| --- | --- | --- | --- | --- | --- | --- | --- | --- | --- | --- |
| rs560887 | 2 | 169,763,148 | T | C | G6PC2/ABCB11 | 0.44 | 0.10 | 0.11 | 0.52 | **2.6 x10<sup>-13</sup>** |
| rs7713317 | 5 | 95,716,722 | G | A | AC104123.1 | 0.98 | 0.91 | 0.43 | **1.4 x10<sup>-13</sup>** | 1.6 x10<sup>-11</sup> |
| rs2206734 | 6 | 20,694,884 | T | C | CDKAL1 | 0.080 | 0.016 | **6.4 x10<sup>-10</sup>** | 0.46 | 1.4 x10<sup>-9</sup> |
| rs11558471 | 8 | 118,185,733 | G | A | SLC30A8 | 0.71 | 0.21 | 5.8 x10<sup>-4</sup> | 4.2 x10<sup>-13</sup> | **4.8 x10<sup>-15</sup>** |
| rs11187144 | 10 | 94,469,980 | C | T | HHEX | 0.91 | 0.88 | **2.7 x10<sup>-8</sup>** | **0.81** | 2.9 x10<sup>-6</sup> |
| rs7903146 | 10 | 114,758,349 | T | C | TCF7L2 | 0.034 | 0.0046 | 0.0023 | 3.5 x10<sup>-18</sup> | **4.2 x10<sup>-20</sup>** |
| rs11038913 | 11 | 46,559,730 | C | T | AMBRA1 | 0.071 | 0.028 | 0.12 | **4.9 x10<sup>-18</sup>** | 1.6 x10<sup>-17</sup> |
| rs11039182 | 11 | 47,346,723 | C | T | MADD | 0.81 | 0.55 | - | **3.4 x10<sup>-45</sup>** | 6.8 x10<sup>-44</sup> |
| rs1552224 | 11 | 72,433,098 | C | A | ARAP1 | 0.085 | 0.015 | 0.097 | 3.4 x10<sup>-38</sup> | **1.8 x10<sup>-38</sup>** |
| rs10830956 | 11 | 92,681,013 | T | C | SNRPGP16 | 0.075 | 0.71 | 5.6 x10<sup>-12</sup> | 0.0033 | **1.2 x10<sup>-21</sup>** |
| rs4502156 | 15 | 62,383,155 | C | T | NPM1P47 | 0.65 | 0.67 | 5.4 x10<sup>-6</sup> | 3.8 x10<sup>-11</sup> | **7.4 x10<sup>-15</sup>** |
| rs4790333 | 17 | 2,262,703 | T | C | SGSM2 | 0.023 | 0.040 | 0.37 | **2.2 x10<sup>-8</sup>** | 1.5 x10<sup>-7</sup> |

<sup>a</sup> For each significant region we report *p*-values for the most associated SNP across all five test (the joint test and the four univariate tests). The most significant test per SNP is indicated by a *p*-value in bold.
Missing values are coded as “-”

#### Example 2

In this example, we assume an investigator wants to analyze jointly many phenotypes while assuming a strong a priori on the expected multitrait association pattern. Here we used a set of 20 phenotypes related to metabolism and displaying substantial genetic correlation (**Figure S1**). We applied the *sumZ* test from the command line version of JASS while using the weights based on the loading of the top 10 principal components (PC) of the genetic correlation matrix,), as proposed in the HIPO method^10^. Genetic and null correlation were derived using the *LDscore* approach^6^. Each of the 10 analyses took less than 3mn, illustrating the strong usability of JASS for a fast exploration of various alternative multivariate tests. For illustrating purposes, we present in **Figure 5** the quadrant plot resulting from the analysis based on PC1.

**Figure 5:**
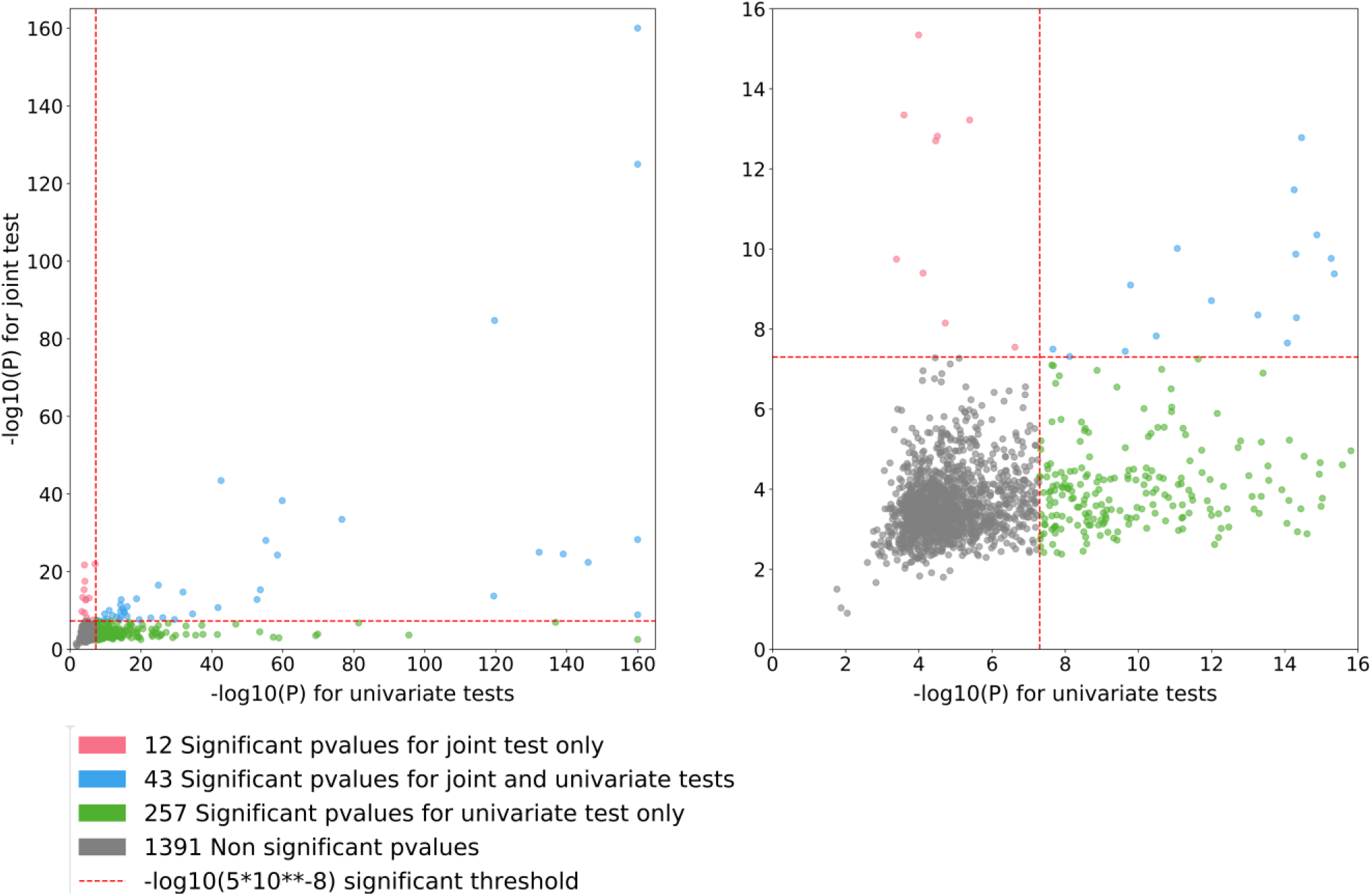
Quadrant plots derived from example 2. The quadrant plot shows the best signal per region from the multivariate test (y-axis) as a function of the best signal for the same region from the univariate analysis (*x*-axis). We focused on the sumZ test using weights defined as the loadings of the first principal component of the genetic correlation matrix times the inverse of the covariance matrix. Green dots represent regions identified by the univariate test only, red dots represent regions identified by the multivariate test only, and blue dots are regions identified by both approaches. Left panel includes all regions (note that −log10(p-value) larger than 160 have been replace by 160). Right panel is a zoom centered around the genome-wide significance level (*p*=5×10^−8^).

The univariate GWAS identified a total of 300 associated loci. Overall, each single PC analysis detected a few additional loci while missing approximately two third of those identified by the univariate screenings. As shown in **Figure 6a** for the first three PCs, the overlap across each PC was relatively strong. We summarized in **Figure 6b** the detection of new loci identified for each PC, and the cumulative number when using an increasing number of PCs. With 10 PCs, the total number of new loci equals 101 if using the standard 5×10^−8^ *p*-value threshold, or 67 if using a more stringent threshold accounting for the 10 tests performed (i.e. 5×10^−9^).

**Figure 6:**
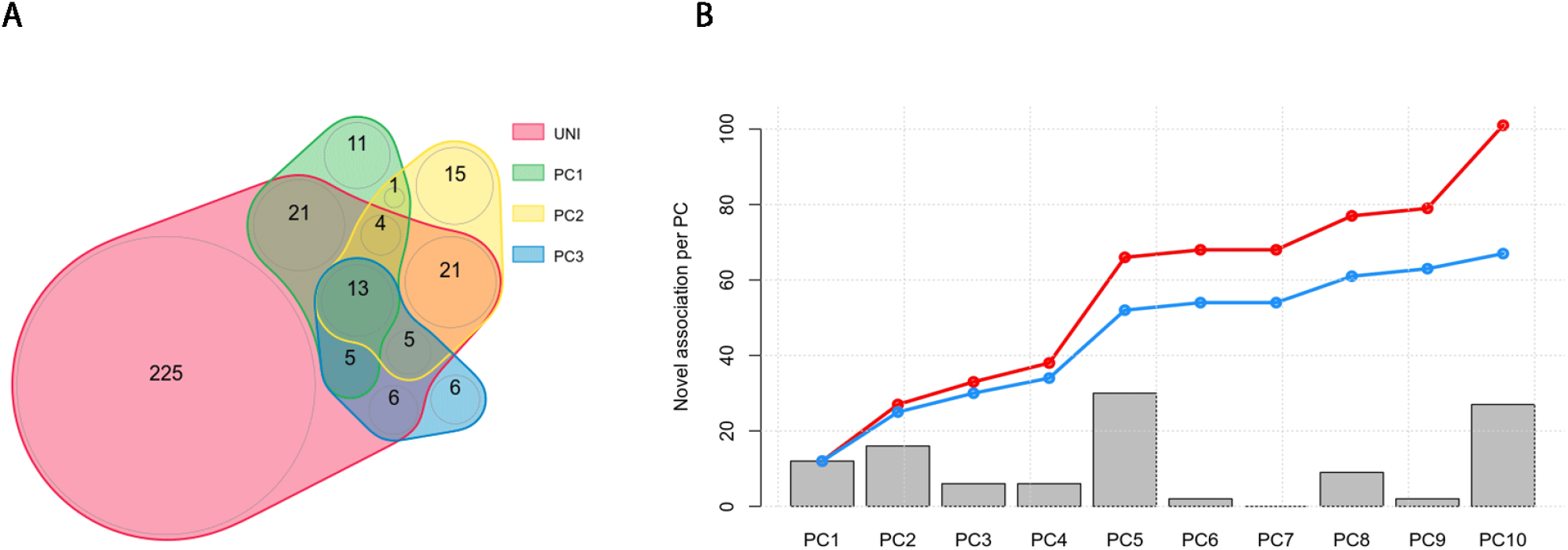
Overview of results from example 2. We performed the *sumZ* test for the analysis of 20 phenotypes while using the weights proposed in the HIPO approach −i.e. using the loadings from the 10 first principal components (PC) of the genetic correlation matrix, weighted by the inverse of the covariance matrix. **Panel A** shows the overlap of identified loci across the univariate screeening and the top 3 PCs. **Panel B** shows the number of loci found associated for each PC on top of those identified by the univariate screening at a *p*-value threshold of 5×10^−8^. The red line indicates the cumulative number of additional signal when merging new signals from an increasing number of PCs. The blue line indicates the same cumulative number of new signal but after applying Bonferroni correction accounting for the total number of PCs analyzed.

#### Example 3

In the last example, we consider a scenario where an investigator wants to confirm the relevance of SNPs near genome-wide significance for Crohn’s diseases (CD) through in-silico replication of association across three other inflammatory conditions: ulcerative colitis (UC), rheumatoid arthritis (RA), and asthma. We show that performing multitrait analysis on these phenotypes related to the primary outcome can improve the validation of these variants. All analyses were performed using summary statistics from the aforementioned traits described in **Table S4**. In practice, we first extracted for CD the most associated SNP for each of the 1704 regions from Berisa and Pickrell and classified those top SNPs in three categories: i) those with *p*-value below genome-wide significance, ii) those suggestive for significance (i.e. having a *p*-value between 1×10^−6^ and 5×10^−8^), and those not significant. There were 33 candidate suggestive significant SNPs (group ii, **Table S5**) for our replication analysis. For each of these SNPs we extracted the *p*-value for association for the three phenotypes and from the omnibus test derived using the command line version of JASS.

Of the 33 SNPs, 24% (N=8), 18% (N=6), and 61% (N=20) were replicated at the Bonferroni corrected *p*-value threshold of 0.0015 (i.e. 0.05/33), for Asthma, RA, and UC, respectively. The omnibus test outperformed all individual univariate signals with an overall replication of 76% (N=25), while performing a single test instead of three. Overall, 7 out of the 33 SNPs had *p*-values below the genome-wide significance threshold (*P*<5×10^−8^) with at least one of the approach (**Table 2**). The omnibus test highlighted in particular two SNPs not identified by the three individual GWAS. The first one, rs3184504, is a missense mutation in *SH2B3.* This variant is cited in more than 40 publications, many of them related to T1D, celiac disease and other autoimmune disorder. The second variant, rs267949, is an intron variant of *DAP* on chromosome 5. While not identified in the univariate GWAS of we used, a previous study found association between the *DAP* gene and UC^33^.

**Table 2:** Validation of SNPs from example 2

| RSID | Chr. | Position | Ref. | Alt. | Gene | P<sub>CD</sub> | P<sub>Asthma</sub> | P<sub>RA</sub> | P<sub>UC</sub> | P<sub>JASS</sub> |
| --- | --- | --- | --- | --- | --- | --- | --- | --- | --- | --- |
| rs7514098 | 1 | 8,177,632 | A | G | LOC107984915 | 9.8 x10<sup>-8</sup> | 0.64 | 0.027 | **4.1 x10<sup>-11</sup>** | **3.3 x10<sup>-9</sup>** |
| rs4845604 | 1 | 151,801,680 | A | G | RORC | 6.1 x10<sup>-7</sup> | 0.16 | 0.37 | **1.6 x10<sup>-11</sup>** | **2.3 x10<sup>-9</sup>** |
| rs267949 | 5 | 10,743,929 | T | C | DAP | 1.3 x10<sup>-7</sup> | 0.063 | 9.3 x10<sup>-7</sup> | 4.9 x10<sup>-5</sup> | **1.5 x10<sup>-8</sup>** |
| rs174564 | 11 | 61,588,305 | G | A | FADS2 | 7.1 x10<sup>-7</sup> | **1.4 x10<sup>-8</sup>** | 5.7 x10<sup>-4</sup> | 0.061 | **6.0 x10<sup>-10</sup>** |
| rs3184504 | 12 | 111,884,608 | T | C | SH2B3 | 5.7 x10<sup>-7</sup> | 8.5 x10<sup>-5</sup> | 3.0 x10<sup>-7</sup> | 5.5 x10<sup>-6</sup> | **1.2 x10<sup>-12</sup>** |
| rs2836881 | 21 | 40,466,299 | T | G | LOC107985484 | 5.7 x10<sup>-7</sup> | 0.31 | 0.41 | **1.1 x10<sup>-32</sup>** | **3.5 x10<sup>-27</sup>** |
| rs138788 | 22 | 35,729,721 | G | A | TOM1 | 7.2 x10<sup>-8</sup> | 0.14 | 0.58 | **2.9 x10<sup>-8</sup>** | 1.4 x10<sup>-6</sup> |

## Discussion

The past few years saw a dramatic increase in the number of publicly available GWAS summary statistics for a broad range of phenotypes. This wealth of data is coming along a strong interest from the community for multitrait analysis, and mulvariate association testing in particular. In this study, we present JASS, a command line and web-based package dedicated to the joint analysis of GWAS summary statistics. JASS addresses the need for a fast and user-friendly tool to perform various joint analysis of summary statistics. Our package includes the two most popular multivariate approaches, an omnibus test and a weighted sum of Z-scores, but allows for alternative approaches to be implemented by advanced users. JASS also includes several complementary functions for both a dynamic synthesis and visualization of large-scale results, and for the pre-processing of heterogeneous GWAS data, a critical step for valid multivariate analysis.

Using existing GWAS summary statistics, we showed the flexibility and strong computational performances of our package as compared to the existent. We further performed three arbitrary real data applications to demonstrate the potential of JASS. These examples cover various scenarios, including the joint analysis of related phenotypes for the identification of new associated loci missed by univariate analyses, the exploratory analysis of alternative weighting schemes to identify specific genetic components across a large number of traits, and the validation of suggestive associations for a given phenotype using multivariate analysis applied to correlated phenotypes. We also deployed a publicly available web instance of JASS (http://jass.pasteur.fr) currently including 49 pre-processed and harmonized GWAS summary statistics, and covering several common diseases and quantitative traits. This installation-free instance of JASS allows non-expert to performed various complementary analyses relevant to their specific study.

There are various multivariate analyses that can be performed from a given set of of GWAS summary statistics. Different tests (e.g. *omnibus* or *sumZ,* as implemented in JASS), different parameters (e.g. alternative weighting scheme for *sumZ),* and the choice of a subset of phenotypes to be analyzed jointly, will lead to the identification of different loci. To our knowledge there are no established guidelines for setting an optimal approach. Moreover, as discussed in previous studies, alternative models likely capture complementary compononents of the genetic architecture of the traits under study. The JASS packages not only offers the possibility to explore quickly a range of alternative models, but it is also a first step toward building an integrated platform including both multitrait association testing and the generation of biological hypothesis on the underlying genetic structure.

## Data Availability

A JASS public server currently including 49 clean and harmonized GWAS is available at http://jass.pasteur.fr For local use, the source code of JASS can be found at https://gitlab.pasteur.fr/statistical-genetics/jass. Installation, configuration and data import instructions are included and linked from the README.md file. The JASS software is released under the terms of the MIT license (see https://gitlab.pasteur.fr/statistical-genetics/jass/blob/master/LICENSE). The pre-processing package is available here https://gitlab.pasteur.fr/statistical-genetics/JASS_Pre-processing.

## Supplementary Data

Supplementary Data include five tables and one figure.

## Acknowledgments

The authors wish to thank Bertrand Néron and Thomas Cockelaer for their useful advice on *Python* software development, as well as Emmanuel Guichard and the production team of the Institut Pasteur IT department for providing the infrastructure to deploy the JASS public server.

## References

1. Buniello, A., MacArthur, J.A.L., Cerezo, M., Harris, L.W., Hayhurst, J., Malangone, C., McMahon, A., Morales, J., Mountjoy, E., Sollis, E. et al. (2019) The NHGRI-EBI GWAS Catalog of published genome-wide association studies, targeted arrays and summary statistics 2019. Nucleic Acids Res, 47, D1005–D1012.

2. Pasaniuc, B. and Price, A.L. (2017) Dissecting the genetics of complex traits using summary association statistics. Nat Rev Genet, 18, 117–127.

3. Turley, P., Walters, R.K., Maghzian, O., Okbay, A., Lee, J.J., Fontana, M.A., Nguyen-Viet, T.A., Wedow, R., Zacher, M., Furlotte, N.A. et al. (2018) Multi-trait analysis of genome-wide association summary statistics using MTAG. Nat Genet, 50, 229–237.

4. Pasaniuc, B., Zaitlen, N., Shi, H., Bhatia, G., Gusev, A., Pickrell, J., Hirschhorn, J., Strachan, D.P., Patterson, N. and Price, A.L. (2014) Fast and accurate imputation of summary statistics enhances evidence of functional enrichment. Bioinformatics, 30, 2906–2914.

5. Kichaev, G., Yang, W.Y., Lindstrom, S., Hormozdiari, F., Eskin, E., Price, A.L., Kraft, P. and Pasaniuc, B. (2014) Integrating functional data to prioritize causal variants in statistical fine-mapping studies. PLoS Genet, 10, e1004722.

6. Bulik-Sullivan, B., Finucane, H.K., Anttila, V., Gusev, A., Day, F.R., Loh, P.R., ReproGen, C., Psychiatric Genomics, C., Genetic Consortium for Anorexia Nervosa of the Wellcome Trust Case Control, C., Duncan, L. et al. (2015) An atlas of genetic correlations across human diseases and traits. Nat Genet, 47, 1236–1241.

7. Liu, Z. and Lin, X. (2018) Multiple phenotype association tests using summary statistics in genome-wide association studies. Biometrics, 74, 165–175.

8. Cichonska, A., Rousu, J., Marttinen, P., Kangas, A.J., Soininen, P., Lehtimaki, T., Raitakari, O.T., Jarvelin, M.R., Salomaa, V., Ala-Korpela, M. et al. (2016) metaCCA: summary statistics-based multivariate metaanalysis of genome-wide association studies using canonical correlation analysis. Bioinformatics, 32, 1981–1989.

9. Qi, G. and Chatterjee, N. (2018) Heritability informed power optimization (HIPO) leads to enhanced detection of genetic associations across multiple traits. PLoS Genet, 14, e1007549.

10. Wang, Z., Sha, Q. and Zhang, S. (2016) Joint Analysis of Multiple Traits Using “Optimal” Maximum Heritability Test. PLoS One, 11, e0150975.

11. Zhu, X., Feng, T., Tayo, B.O., Liang, J., Young, J.H., Franceschini, N., Smith, J.A., Yanek, L.R., Sun, Y.V., Edwards, T.L. et al. (2015) Meta-analysis of correlated traits via summary statistics from GWASs with an application in hypertension. Am J Hum Genet, 96, 21–36.

12. Kim, J., Bai, Y. and Pan, W. (2015) An Adaptive Association Test for Multiple Phenotypes with GWAS Summary Statistics. Genet Epidemiol, 39, 651–663.

13. Province, M.A. and Borecki, I.B. (2013) A correlated meta-analysis strategy for data mining “OMIC” scans. Pac Symp Biocomput 236–246.

14. Ray, D. and Boehnke, M. (2018) Methods for meta-analysis of multiple traits using GWAS summary statistics. Genet Epidemiol, 42, 134–145.

15. van der Sluis, S., Posthuma, D. and Dolan, C.V. (2013) TATES: efficient multivariate genotype-phenotype analysis for genome-wide association studies. PLoS Genet, 9, e1003235.

16. O’Reilly, P.F., Hoggart, C.J., Pomyen, Y., Calboli, F.C., Elliott, P., Jarvelin, M.R. and Coin, L.J. (2012) MultiPhen: joint model of multiple phenotypes can increase discovery in GWAS. PLoS One, 7, e34861.

17. Zhou, X. and Stephens, M. (2014) Efficient multivariate linear mixed model algorithms for genome-wide association studies. Nat Methods, 11, 407–409.

18. Aschard, H., Vilhjalmsson, B.J., Greliche, N., Morange, P.E., Tregouet, D.A. and Kraft, P. (2014) Maximizing the power of principal-component analysis of correlated phenotypes in genome-wide association studies. Am J Hum Genet, 94, 662–676.

19. Geihs, M., Yan, Y., Walter, K., Huang, J., Memari, Y., Min, J.L., Mead, D., Consortium, U.K., Hubbard, T.J., Timpson, N.J. et al. (2015) An interactive genome browser of association results from the UK10K cohorts project. Bioinformatics, 31, 4029–4031.

20. Juliusdottir, T., Banasik, K., Robertson, N.R., Mott, R. and McCarthy, M.I. (2018) Toppar: an interactive browser for viewing association study results. Bioinformatics, 34, 1922–1924.

21. Pruim, R.J., Welch, R.P., Sanna, S., Teslovich, T.M., Chines, P.S., Gliedt, T.P., Boehnke, M., Abecasis, G.R. and Willer, C.J. (2010) LocusZoom: regional visualization of genome-wide association scan results. Bioinformatics, 26, 2336–2337.

22. Khramtsova, E.A. and Stranger, B.E. (2017) Assocplots: a Python package for static and interactive visualization of multiple-group GWAS results. Bioinformatics, 33, 432–434.

23. Liu, Z. and Lin, X. (2018) Multiple phenotype association tests using summary statistics in genome-wide association studies. Biometrics, 74, 165–175.

24. Berisa, T. and Pickrell, J.K. (2016) Approximately independent linkage disequilibrium blocks in human populations. Bioinformatics, 32, 283–285.

25. Genomes Project, C., Auton, A., Brooks, L.D., Durbin, R.M., Garrison, E.P., Kang, H.M., Korbel, J.O., Marchini, J.L., McCarthy, S., McVean, G.A. et al. (2015) A global reference for human genetic variation. Nature, 526, 68–74.

26. Julienne, H., Shi, H., Pasaniuc, B. and Aschard, H. (2018) RAISS: Robust and Accurate imputation from Summary Statistics. bioRxiv, 502880.

27. Schizophrenia Working Group of the Psychiatric Genomics, C. (2014) Biological insights from 108 schizophrenia-associated genetic loci. Nature, 511, 421–427.

28. Bycroft, C., Freeman, C., Petkova, D., Band, G., Elliott, L.T., Sharp, K., Motyer, A., Vukcevic, D., Delaneau, O., O’Connell, J. et al. (2018) The UK Biobank resource with deep phenotyping and genomic data. Nature, 562, 203–209.

29. Prive, F., Aschard, H., Ziyatdinov, A. and Blum, M.G.B. (2018) Efficient analysis of large-scale genome-wide data with two R packages: bigstatsr and bigsnpr. Bioinformatics, 34, 2781–2787.

30. Yang, G., Sau, C., Lai, W., Cichon, J. and Li, W. (2015) USAT: A Unified Score-based Association Test for Multiple Phenotype-Genotype Analysis. 344, 1173–1178.

31. Bouatia-Naji, N., Rocheleau, G., Van Lommel, L., Lemaire, K., Schuit, F., Cavalcanti-Proenca, C., Marchand, M., Hartikainen, A.L., Sovio, U., De Graeve, F. et al. (2008) A polymorphism within the G6PC2 gene is associated with fasting plasma glucose levels. Science, 320, 1085–1088.

32. Rose, C.S., Grarup, N., Krarup, N.T., Poulsen, P., Wegner, L., Nielsen, T., Banasik, K., Faerch, K., Andersen, G., Albrechtsen, A. et al. (2009) A variant in the G6PC2/ABCB11 locus is associated with increased fasting plasma glucose, increased basal hepatic glucose production and increased insulin release after oral and intravenous glucose loads. Diabetologia, 52, 2122–2129.

33. Anderson, C.A., Boucher, G., Lees, C.W., Franke, A., D’Amato, M., Taylor, K.D., Lee, J.C., Goyette, P., Imielinski, M., Latiano, A. et al. (2011) Meta-analysis identifies 29 additional ulcerative colitis risk loci, increasing the number of confirmed associations to 47. Nat Genet, 43, 246–252.

