## Supplementary materials for "JASS: Command Line and Web interface for the joint analysis of GWAS results"

### Supplementary data

### Table S1. JASS command line options

| command | description | Arguments |
|---|---|---|
| *Jass()* | Import pre-processed data | --input-data-path *"PREPROC/*.txt"*<br>--init-covariance-path *"Covariance_final.csv"*<br>--regions-map-path *"regions.txt"*<br>--description-file-path *"description_metabo.txt"*<br>--init-table-path *"initTable.hdf5"* |
| *create-inittable()* | Create the iniTable.hdf5 files | --phenotypes *GWAS1 GWAS2 GWAS3*<br>--worktable-path *workTable.hdf5* |
| *list-phenotypes()* | List available phenotypes | |
| *create-projectdata()* | Compute the joint statistics and generate plots | |
| *create-worktable()* | Compute joint statistics for a given set of GWAS | |
| *plot-manhattan()* | Generate Manhattan plot | |
| *plot-quadrant()* | Generate a quadrant plot | |
| *serve()* | Run JASS web server | |

### Table S2: features of the three version of JASS

| Feature | Public Pasteur server (jass.pasteur.fr) | Local, web server access | Local, command line |
|---|---|---|---|
| Customize covariance matrix | No | Not so easy | Yes |
| Custom regions | No | Not so easy | Yes |
| Dynamic plots | Yes | Yes | No |
| Works on imputed data | No | Maybe | Yes |
| Add own GWAS results | No | Yes | Yes |
| Use SumZ | No | No | Yes |
| Custom integration statistic | No | No | Yes |

### Table S3: Other multi-GWAS methods functionality

| Method | Language | Code availability | Documentation | Harmonizing tool | Command line | Web interface | Missing values |
|---|---|---|---|---|---|---|---|
| JASS | Python3 package | https://gitlab.pasteur.fr/statistical-genetics/jass | http://statistical-genetics.pages.pasteur.fr/jass/index.html | Yes (companion script) https://gitlab.pasteur.fr/statistical-genetics/JASS_Pre-processing | Yes | Yes | Yes |
| HIPPO (Qi et al) | R package | https://github.com/gqi/hipo | Github Readme with a tutorial + R man pages | No | No | No | No |
| aSPU (Kim et al) | R package | https://cran.r-project.org/web/packages/aSPU/index.html | Github Readme with a tutorial + R man pages | No | No | No | No |
| MTAG (REF) | Python2.7 script | https://github.com/omeed-maghzian/mtag | Github Readme and wiki | No | Yes | No | No |
| CPASSOC (Zhu et al) | R functions | http://hal.case.edu/~xxz10/zhu-web/ | Word document | No | No | No | No |
| MHT-O (Wang et al) | R functions | http://pages.mtu.edu/~shuzhang/software.html | No | No | No | No | No |
| USAT (Ray et al) | No code* | No | No | No | No | No | No |
| M Omics (Province et al) | No code* | No | No | No | No | No | No |

**No link provided with the manuscript*

### Table S4: GWAS available in the online version of JASS as of 04/2019

| Consortium | Outcome | Description | Reference |
|---|---|---|---|
| CARDIOGRAM-C4D | CHD | Coronary Heart Disease | *Peden et al. 2011* |
| | CHD | Coronary Heart Disease | *Schunkert et al. 2011* |
| | CAD | coronary artery disease | *Nikpay et al 2015* |
| DIAGRAM | T2D | Type 2 Diabetes | *Morris et al. 2012* |
| GEFOS | BMD-FOREARM | Bone Mineral Density Forearm | *Zheng et al. 2015* |
| | BMD-NECK | Bone Mineral Density Neck | |
| | BMD-SPINE | Bone Mineral Density Spine | |
| GIANT | BMI | Body Mass Index | *Locke et al* |
| | HIP | Hip Circumference | *Shungin et al. 2015* |
| | WHR | Waist hip ratio | |
| | WC | Waist Circumference | |
| | HEIGHT | Height | *Wood et al* |
| GLG | HDL | High-density lipoprotein | *Teslovich et al. 2010* |
| | TG | Triglyceride | |
| | TC | Total Cholesterol | |
| | LDL | Low-density lipoprotein | |
| IGAP | AZ | Alzeimer | *Lambert et al. 2013* |
| IMSGC-WTCC2 | MULTSCLE | Multiple sclerosis | *Sawcer et al. 2011* |
| KIDNEY | CKD | Chronic Kidney Disease | *Pattaro et al. 2016* |
| | EGFR-CYS | Glomerular Filtration Rate from Cystatin C | |
| | EGFR-CREA | Glomerular Filtration Rate from Creatinine | |
| LOOS | BEATSTMN | beats/min | *den Hoed et al. 2013* |
| MAGIC | HOMA-IR | Insulin Resistance | *Dupuis et al. 2010* |
| | FAST-INSULIN | Fasting Insulin | |
| | FAST-GLUCOSE | Fasting Glucose | |
| | HOMA-B | Beta-cell Function | |
| | IS | Insulin secretion | *Prokopenko et al. 2014* |
| | 2HGLU-ADJBMI | 2h Glucose Adjusted For BMI | *Saxena et al. 2010* |
| | HBA1C | Haemoglobin A1c | *Soranzo et al. 2010* |
| | FPI | Fasting proinsulin | *Strawbridge et al. 2011* |
| MEGASTROKE | AS | any stroke | *Malik et al. 2018* |
| | CES | cardioembolic stroke | |
| | AIS | any ischemic stroke | |
| NEIGHBORHOOD | POAG | Primary Open Angle Glaucoma | *Bailey et al. 2016* |
| PGC | MDD | Major Depressive Disorder | *Ripke et al* |
| | SCZ | Schizophrenia | |
| | BIP | Bipolar Disorder | *Sklar et al. 2011* |
| RA | RA | Rheumatoid Arthritis | *Okada et al. 2014* |
| ReproGen | AME | Age at Menopause | *Day et al* |
| SSGAC | COLLEGE | College Completion | *Rietveld et al. 2013* |
| | EDUYEAR | Years of Educational Attainment | *Rietveld et al. 2013* |
| TAGC | ASTHMA | Asthma | *Demenais et al. 2018* |
| UKIBD | IBD | inflammatory bowl disease | *deLange et al. 2017* |
| | UC | ulcerative colitis | |
| | CD | Crohn | |
| UKPSC | CHOLANG | Primary sclerosing cholangitis | *Ji et al. 2017* |
| VAN-HEEL | CELIAC | Celiac disease | *Dubois et al. 2010* |
| VGHRV | SDNN | standard deviation of the normal-to-normal interval | *Nolte et al. 2017* |
| WILLER | ATRIALFIBRI | Atrial Fibrillation | *Nielsen et al. 2018* |

### Table S5: Detailed results from example 3

| RSID | Chr. | position | Ref. | Alt. | PCD | Pasthma | Pra | Puc | Pjass |
|---|---|---|---|---|---|---|---|---|---|
| rs7514098 | 1 | 8,177,632 | A | G | 9.8E-8 | 0.64 | 0.027 | **4.1E-11** | **3.3E-9** |
| rs111471231 | 1 | 38,894,402 | T | C | 4.8E-7 | 0.058 | 0.22 | 9.1E-3 | 0.011 |
| rs4845604 | 1 | 151,801,680 | A | G | 6.1E-7 | 0.16 | 0.37 | **1.6E-11** | **2.3E-9** |
| rs7526313 | 1 | 209,958,599 | C | T | 7.1E-7 | 0.056 | 0.15 | **5.3E-7** | **3.8E-6** |
| rs11901880 | 2 | 141,605,983 | G | A | 7.0E-7 | 0.017 | 0.32 | **8.7E-4** | **7.4E-4** |
| rs57669318 | 2 | 198,823,642 | A | C | 6.0E-8 | **9.0E-5** | **3.7E-5** | 0.11 | **2.2E-7** |
| rs13001905 | 2 | 242,476,987 | G | A | 8.9E-7 | 0.35 | 9.5E-3 | **1.2E-4** | **8.2E-5** |
| rs12511665 | 4 | 48,213,214 | T | C | 3.7E-7 | 0.11 | **3.8E-6** | 0.75 | **2.7E-5** |
| rs267949 | 5 | 10,743,929 | T | C | 1.3E-7 | 0.063 | **9.3E-7** | **4.9E-5** | **1.5E-8** |
| rs27460 | 5 | 111,010,157 | G | A | 6.6E-7 | 0.36 | 0.39 | **7.9E-5** | 1.9E-3 |
| rs2428669 | 5 | 128,295,386 | A | G | 1.1E-7 | 1.9E-3 | 0.047 | 0.018 | **4.0E-4** |
| rs1267500 | 6 | 14,715,825 | C | T | 9.1E-8 | 0.075 | 0.68 | **4.8E-7** | **1.3E-5** |
| rs9369329 | 6 | 42,029,905 | T | C | 2.0E-7 | **1.1E-4** | 1.00 | **1.1E-3** | **4.1E-5** |
| rs206203 | 7 | 20,374,252 | A | C | 7.4E-7 | 3.7E-3 | 1.00 | **2.0E-4** | **1.1E-4** |
| rs243514 | 7 | 148,427,873 | A | G | 5.2E-7 | 0.22 | 0.71 | **2.3E-5** | **4.5E-4** |
| rs10974513 | 9 | 449,333 | A | G | 5.0E-7 | 0.51 | 0.20 | **2.3E-5** | **3.4E-4** |
| rs10815099 | 9 | 4,876,202 | C | T | 3.0E-7 | 0.021 | 0.77 | **1.0E-3** | 2.6E-3 |
| rs7894089 | 10 | 126,419,153 | C | T | 3.8E-7 | 0.66 | 0.71 | **9.2E-7** | **8.4E-5** |
| rs7952451 | 11 | 774,715 | G | A | 1.1E-7 | 0.42 | 0.32 | **1.4E-4** | 2.8E-3 |
| rs174564 | 11 | 61,588,305 | G | A | 7.1E-7 | **1.4E-8** | **5.7E-4** | 0.061 | **6.0E-10** |
| rs593982 | 11 | 65,513,107 | T | C | 3.7E-7 | **1.9E-4** | 0.11 | 1.7E-3 | **3.6E-5** |
| rs12578573 | 12 | 6,842,959 | T | C | 2.0E-7 | 0.71 | 0.20 | 4.8E-3 | 0.028 |
| rs3184504 | 12 | 111,884,608 | T | C | 5.7E-7 | **8.5E-5** | **3.0E-7** | **5.5E-6** | **1.2E-12** |
| rs9549249 | 13 | 41,224,442 | T | C | 1.4E-7 | **2.4E-6** | 0.026 | 0.024 | **2.0E-6** |
| rs12878219 | 14 | 35,381,137 | A | G | 5.5E-7 | 0.70 | 5.4E-4 | 0.039 | **9.9E-4** |
| rs1136165 | 14 | 103,988,180 | G | T | 3.9E-7 | 0.078 | 0.41 | 3.5E-3 | 0.014 |
| rs7220929 | 17 | 80,801,483 | G | A | 7.1E-7 | 0.85 | 1.00 | **2.1E-4** | 6.5E-3 |
| rs2074621 | 19 | 15,290,412 | G | A | 5.2E-7 | 0.021 | 0.55 | 0.53 | 0.021 |
| rs34378208 | 19 | 49,398,405 | T | C | 3.8E-7 | **8.9E-5** | 0.019 | 0.98 | **2.0E-4** |
| rs62194743 | 20 | 15,781,696 | T | C | 7.7E-8 | **5.1E-5** | 0.72 | 0.27 | **7.0E-4** |
| rs113989515 | 20 | 57,840,396 | T | C | 3.7E-7 | 0.11 | 0.010 | **5.6E-6** | **1.1E-5** |
| rs2836881 | 21 | 40,466,299 | T | G | 5.7E-7 | 0.31 | 0.41 | **1.1E-32** | **3.5E-27** |
| rs138788 | 22 | 35,729,721 | G | A | 7.2E-8 | 0.14 | 0.58 | **3.0E-8** | **1.4E-6** |

**Figure S1: Genetic correlation across phenotypes from example 2**

Genetic correlation between the 20 phenotypes derived using the *LDscore* regression

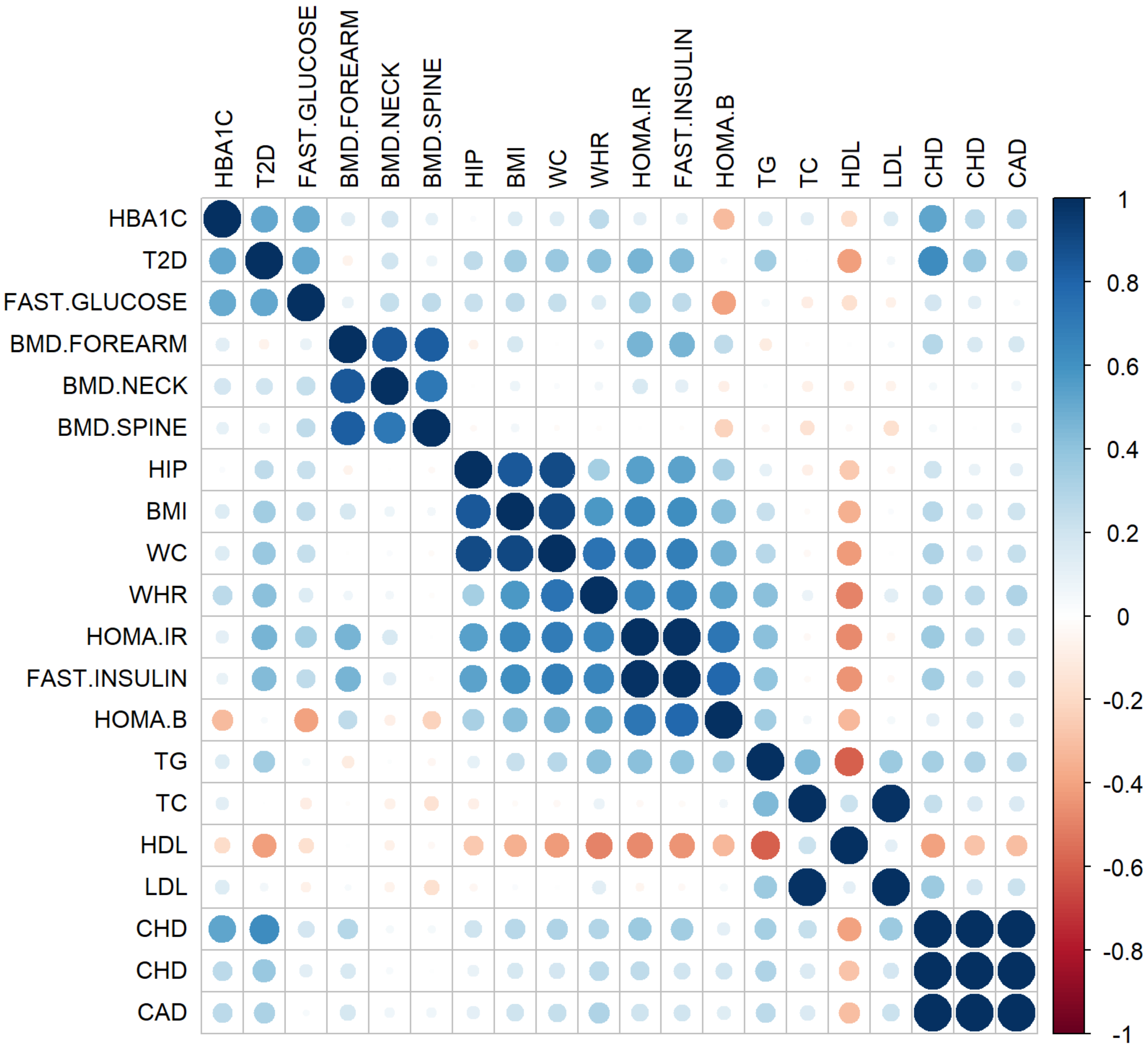
